# Age and cognitive load affect muscle activation profiles in response to physical perturbations while walking

**DOI:** 10.1101/2022.05.29.493879

**Authors:** Uri Rosenblum, Itshak Melzer, Michael Friger, Gabi Zeilig, Meir Plotnik

## Abstract

To maintain balance during walking, the central nervous system must adjust the base of support (i.e., modulation of step length and step width) to the center of mass displacement in every step. We aimed to explore age and concurrent cognitive attention-demanding task effects on lower limb muscle fiber type recruitment in response to unexpected loss of balance during walking i.e., perturbation. Twenty young (YA) and 18 older adults (OA), (27.00±2.79 and 70.13±3.95 years old, respectively) were exposed to unexpected perturbations, while walking on a treadmill, in virtual reality environment. Surface electromyography (sEMG) total spectral power for frequency bands associated with muscle fibers type I (40-60Hz), type IIa (60-150Hz) and type IIb (150-250Hz), from tibialis-anterior and vastus-lateralis muscles were analyzed. Four Generalized Estimating Equations models assessed age and cognitive attention-demanding task’s load association with lower-limb muscle activation patterns resulting from perturbation in single- and double-support phases of the gait cycle. Results show that OA employ a muscle fiber type IIa dominant increase strategy while YA show muscle fiber type IIb dominant increase in muscle fiber type recruitment in response to unexpected perturbations during walking. This suggests that the ability to recruit fast-twitch muscle fibers is deteriorated with age and thus may be related to insufficient balance recovery response.

## Introduction

Studies show that the ability to control balance during walking while performing a concurrent cognitive task (i.e., dual task condition, DT) is reduced[1–3] and deteriorates with age[3,4]. Many falls occur while walking and concurrently performing additional task, such as having a conversation or using a mobile phone[5– 7]. Therefore, it is essential to understand the effects of attentional demands required while being engaged with a concurrent cognitive load on the different aspects of the balance recovery function when balance is lost unexpectedly during walking. Studies show that older adults (OA) exhibit a reduced capacity to walk efficiently[8,9], to resolve simultaneous motor and cognitive tasks (i.e., DT)[10–12], and respond to perturbations[9,13,14]. A concurrent cognitive attentional demand may cause motor-cognitive interference. As seen in past studies, a simultaneous performance of a cognitive and any motor task, i.e., dual task (DT), results in deterioration of performance in one or both tasks, relative to performance of each task separately (single-task performance, ST)[15]. An example, of a motor-cognitive interference paradigm is walking with destabilizing perturbations while performing a concurrent cognitive task[16]. Wittenberg et al. systematic review [16] have demonstrated this for perturbations introduced during walking on treadmill and overground with a variety of concurrent tasks. On the other hand, [17,18] found that balance recovery stepping responses during treadmill walking were not affected by a concurrent arithmetic task (i.e. counting backward by seven) among young and older adults, respectively. Different research groups have examined muscle activation patterns and onset in response to perturbations[19–24]. In a recent study we have observed a significant increase in fast-twitch muscle fibers type IIa recruitment in response to destabilizing perturbations during walking in YA, in time window of two seconds after perturbation[23]. The volume of fast-twitch muscle fibers is decreased with age[25]. It is suggested that changes in muscle architecture cause changes in muscle activation patterns. This led McCrum et al.[26] to investigate the association between type II muscle fiber properties and balance recovery following large perturbations during walking in young and older adults. They found no association.

In fact, the research covering these aspects and specific aspects of falling in the elderly population is still scarce. Thus, there is a need to explore age-related differences in muscle fiber type recruitment and DT performance to better understand mechanisms of balance recovery in response to unexpected perturbations during walking. Therefore, in this study we aimed to investigate muscle recruitment patterns when concurrent cognitive attentional demands are high, and there is a competition for allocating attention and further investigate brain computational capacity.

We employed a similar DT paradigm to the one above, with the aim to characterize the effects of ageing and concurrent cognitive attention-demanding task on lower limb muscle activation pattern i.e., total spectral power as reflected in surface electromyogram (sEMG) signals, in different frequency bands, while the participants responded to unannounced perturbations while walking. We hypothesized that: (1) In line with Schillings et al.[22] and due to the loss of fast-twitch muscle fibers with age, YA and OA will exabit a different muscle fiber type recruitment dominancy in response to unexpected perturbations; (2) With regards to age X concurrent cognitive task interaction we hypothesize that both YA and OA will demonstrate elevated total EMG spectral power after perturbation during perturbed DT as compared to perturbed ST conditions.

## Methods

### Participants

Twenty young and 18 OA participated in this study (Table 1). Exclusion criteria include obesity (i.e., body mass index [kg/m^2^] ≥ 30)[27]; orthopedic conditions affecting gait and balance (e.g., total knee replacement, total hip replacement, ankle sprain, limb fracture); cognitive loss (Mini-Mental State Exam (MMSE) < 24[28] or psychiatric conditions; cardiac conditions (e.g., non-stable ischemic heart disease, congestive heart failure); chronic obstructive pulmonary disease; and neurological diseases associated with loss of balance (e.g., multiple sclerosis, myelopathy). The Institutional Review Board of Sheba Medical Center approved the study protocol (Approval Number 9407-12). All participants signed a written informed consent form prior to entering the study.

**Table 1:** Demographic and physical characteristics represented as Median (Range)

|  | YA (N = 20) | OA (N = 18) |
| --- | --- | --- |
| **Age (years)** | 27.50 (22-33) | 69.00 (65-80) |
| **Sex (F/M)** | 10/10 | 9/9 |
| **Height (m)** | 1.69 (1.51-1.84) | 1.70 (1.57-1.90) |
| **Weight (Kg)\*\*** | 61.00 (38.80-85.00) | 71.00 (56.00-95.00) |
| **BMI (kg/m<sup>2</sup>)\*\*** | 22.25 (17.02-27.75) | 26.02 (18.93-30.32) |
| **Years of education** | 15.50 (12.00-20.00) | 15(10-27) |

Abbreviations: YA=Young Adults; OA=Older Adults; SD=Standard Deviation; BMI=Body Mass Index. Mann-Whitney U test was used to compare between the age groups.
\* P<0.05, \*\* P<0.01.

### Apparatus and settings

Our experimental set-up was described elsewhere in detail[23]. Briefly, participants walked in the CAREN High-End (Motek Medical B.V., Amsterdam, Netherlands) virtual reality system, equipped with a split-belt treadmill mounted on a platform that can move in six degrees of freedom. The platform is positioned in the center of a 360° dome-shaped screen. Medio-lateral unannounced support-surface physical perturbations were introduced every 50-70 seconds to reduce an anticipation effect. These occurred by shifting the platform 15cm for 0.92 seconds, either left or right. Eighteen Vicon (Vicon Motion Systems, Oxford, UK) motion capture cameras placed around the dome’s circumference along with force plates under the treadmill’s belts, allowed for continuous recording of kinetic (ground reaction forces) and kinematic (motion) data for real-time/post-hoc gait cycle phase detection. Perturbation control and data recording were done by two computers that integrated kinetic, kinematic and perturbation data.

### sEMG recording

sEMG data were collected from vastus lateralis (VL) and tibialis anterior (TA) muscles, known to take part in maintaining walking balance[20,29–32]. Electrode type (bipolar hydrogel surface electrodes), inter-electrode distance, orientation, placement along the contracted muscle belly, and evaluation of surface musculature followed the Surface electromyography for the Non-Invasive Assessment of Muscles (SENIAM guidelines) and International Society of electrophysiology and Kinesiology (ISEK) guideline[33,34] maximizing confidence that the EMG signal can be attributed primarily to the named muscles. The ground electrode was placed on spinous process of C7. Before electrode application, participants’ skin was shaved and cleaned with an abrasive pad and alcohol to reduce skin impedance. Finally, electrodes and cables were secured to the skin with tape to reduce movement artifacts. Before recording data sEMG signals were tested for specific movements to control for crosstalk effects. Raw sEMG signals were recorded on a tablet computer (Lenovo ThinkPad) at 2048 Hz using an eego referential amplifier (eemagine Medical Imaging Solutions GmbH, Berlin, Germany), with an input impedance (referential) >1GΩ, and weight < 500g.

#### Study protocol

First Participants acclimated to walking in virtual reality (see [35] for more details). Then, they walked at a comfortable, self-selected speed (using the system’s self-paced mode[36]) while being subjected to random, unannounced perturbations at different phases of the gait cycle. Perturbations varied in direction (right-left) and gait cycle phase presentation (double-support/single-support). Trials were conducted under two cognitive load conditions: *single task* (i.e., only walking with perturbations, ST) and *dual task* (DT), during which the participants were asked to perform arithmetic calculations, adding 15 numbers presented once every 2 seconds (i.e., lasting 28 sec), based on a paced auditory serial addition test [37], while walking with perturbations. Experiments consisted of 4 perturbation trials (Blocks) of approximately 10 minutes with 2 minutes rest in between to minimize fatigue (see Fig. 1 for illustration). Two blocks were performed in ST condition and two in DT condition in random order (24 perturbations in each condition; in DT blocks each perturbation was accompanied by the concurrent arithmetic task). Note: only rightward platform translation perturbations in left single-support or double-support, with the left leading limb, were considered due to their relevance to falls and injury mechanisms[38–41]. In addition, recently we showed that young adults have different muscle pattern activations in response to perturbations in single-and double-support[23].

**Figure 1:**
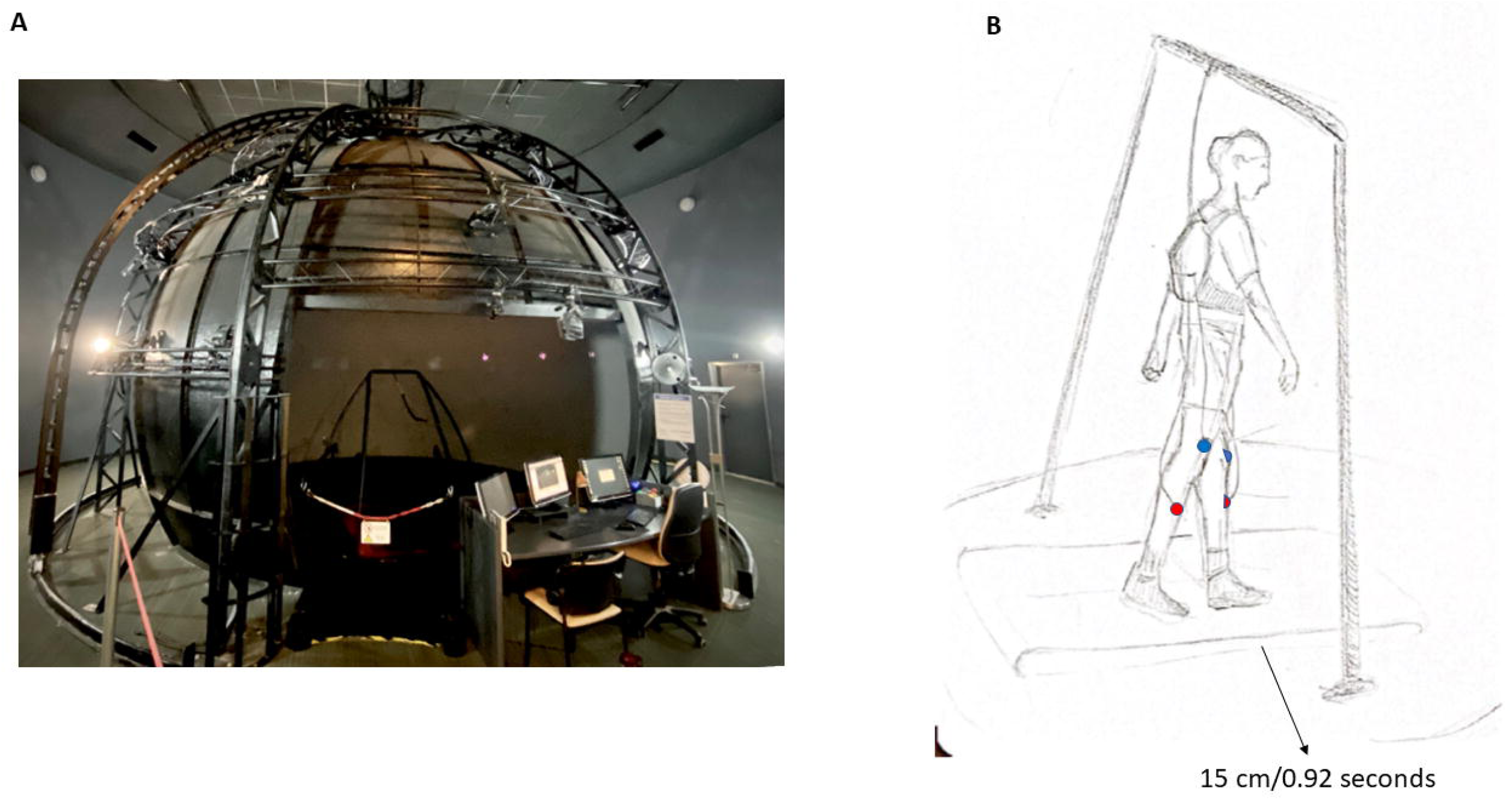
experimental setup. A) CAREN High-End (Motek Medical B.V., Amsterdam, Netherlands) virtual reality system, equipped with a split-belt treadmill mounted on a platform that can move in six degrees of freedom. The platform is positioned in the center of a 360° dome-shaped screen. Eighteen Vicon (Vicon Motion Systems, Oxford, UK) motion capture cameras placed around the dome’s circumference along with force plates under the treadmill’s belts, allowed for continuous recording of kinetic (ground reaction forces) and kinematic (motion) data for real-time/post-hoc gait cycle phase detection. Perturbation control and data recording were done by two computers that integrated kinetic, kinematic and perturbation data. B) Schematic figure of perturbation in double support gait phase. Medio-lateral unannounced support-surface physical perturbations were introduced every 50-70 seconds to reduce an anticipation effect. These occurred by shifting the platform 15cm for 0.92 seconds, either left or right. Raw sEMG signals were recorded on a tablet computer (Lenovo ThinkPad) at 2048 Hz using an eego referential amplifier (eemagine Medical Imaging Solutions GmbH, Berlin, Germany), with an input impedance (referential) >1GΩ, and weight < 500g. These were placed in a backpack on the participants back. sEMG data were collected from vastus lateralis (blue circle) and tibialis anterior (red circle) muscles.

### sEMG data handling, processing, and analyses

sEMG data were preprocessed and analyzed across frequency domains using customized scripts written in MATLAB version R2020b (Mathworks, Nathick USA). Raw sEMG signals were first band-pass filtered (40–1000 Hz) according to the recommendations by Papagiannis et al.,[31] to clean the data from walking and perturbation moving artifact residing in the frequencies ≤ 40 Hz[31,42]. Filtered sEMG signals were sliced in 1-minute windows around perturbation events (i.e., 30 seconds prior to 30 seconds after) and grouped as baseline data, 1 second of undisturbed walking segment, 2 seconds prior to the perturbation, and 2 seconds of balance and walking recovery after perturbation (Seconds-1, & 2, respectively).

sEMG data processing consisted of three main steps: (1) we performed wavelet analysis, a method suitable for non-stationary signals, on the 3 seconds segment (1 seconds baseline + 2 seconds after perturbation) for every perturbation separately; (2) we calculated average spectral power for the following frequency bands: 40-60Hz, 60-150Hz, 150-250Hz, 250-400Hz, and 400-1000Hz; (3) To specifically measure rate and magnitude of change across different frequency bands over time, we calculated a ratio between the total spectral power after the perturbation and baseline, for every frequency band, in every segment where, *fb* is the frequency band in every segment *x* (Second-1 & 2) and *bl* is for the baseline segment:

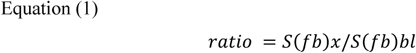

For this study, we examined results for the left and right TA (ankle stabilizing muscle of the non-dominant limb) and VL (anti-gravity muscle of the non-dominant limb), in response to right platform translation during stance of the left leg i.e., before right foot initial contact.

### Dual task accuracy and effect

Dual task accuracy is a performance measure of the cognitive dual task, and was calculated per participant using the following:

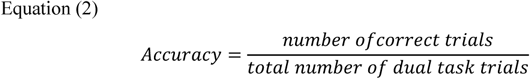

Dual task cost (DTC) represents the change in the total spectral power measure ratio (Equation (1)) consequential of the concurrent cognitive DT, and was calculated per participant, per time bin (i.e., for Seconds-1&2), using the following:

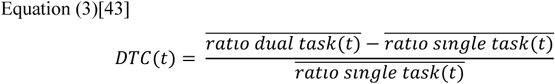

where t stands for the time in seconds (1-2) and ratio is the quotient of equation (3); the median ratio for every participant was used. Positive DTC values represent interference, i.e., increased muscle activation associated with the presence of the cognitive task, and negative values represent facilitation[44,45], i.e., reduced muscle activation.

### Statistical analyses

Data were analyzed using MATLAB version R2020b and SPSS version 25 (IBM Inc., Chicago, IL). The Shapiro-Wilk statistic was first calculated to test for normality of distributions. Groups were compared using Mann–Whitney U test and a chi-square test for between-sex comparisons. To test age-related differences in concurrent cognitive performance, i.e., DT, we conducted a Mann-Whitney U-test to compare concurrent cognitive task accuracy. We used data from 19 YA and 10 OA as one YA and 8 OA had missing cognitive performance data and were excluded from this analysis.

To test our two hypotheses (between age groups and between cognitive load conditions spectral power differences), we performed four GEE models to account for dependence of repeated observations within participants, one for each of the perturbation types (i.e., single-and double-support) and age combinations, to show the age effect more clearly. The change ratio was the independent variable, and time segments (i.e., 2 seconds after the perturbation, Second-1& 2), muscles (i.e., TA and VL), cognitive load conditions (i.e., ST/DT), and frequency bands (i.e., 40-60Hz, 60-150Hz and 150-250Hz) were the within subject variables. Regarding the effect of concurrent cognitive task, the results will be presented in four subgroups: 1) perturbed ST condition in single-support phase; 2) perturbed DT condition in single-support phase; 3) perturbed ST condition in double-support phase; and 4) perturbed DT condition in double-support phase.

The dependent variable ratio (i.e., the total power spectral density after perturbation/baseline) followed a gamma distribution for perturbations during single-support and double-support gait phase. In addition, we transformed the frequency band variables in to three levels (40-60Hz, 60-150Hz, and 150-250Hz), corresponding with Type I, Type IIa and Type IIb related frequencies, respectively [46]. Finally, we focused our analysis on the first two seconds after perturbation in accordance with our previous study, showing the greatest effect of change in total spectral power occurs in this time frame [23]. To quantify the direct effect of a concurrent cognitive task on muscle activation in response to perturbations we calculated the DTC from the EMG signals. Then we used it as the independent variable in four Generalized Estimating Equations (GEE) models for normally distributed dependent variable, similar to the above models to explore the age-related effect for the different muscles i.e., TA and VL, in the first two seconds after perturbation, for YA and OA. Muscles (i.e., TA and VL) and time (i.e., the first two seconds after perturbation) were the within subject variables. Two OA had DTC scores greater than three standard deviations from the mean. They were excluded from the GEE analyses. After excluding the outliers, the DTC followed a normal distribution (i.e., Shapiro-Wilks test p=0.07). Bonferroni correction was applied to account for multiple comparisons. Statistical significance was set *a priori* at p<0.05.

## Results

### Demographic characteristics

### Cognitive performance

OA’s made significantly more cognitive task errors compared to YA (p=0.05, Fig. 2).

**Fig. 2:**
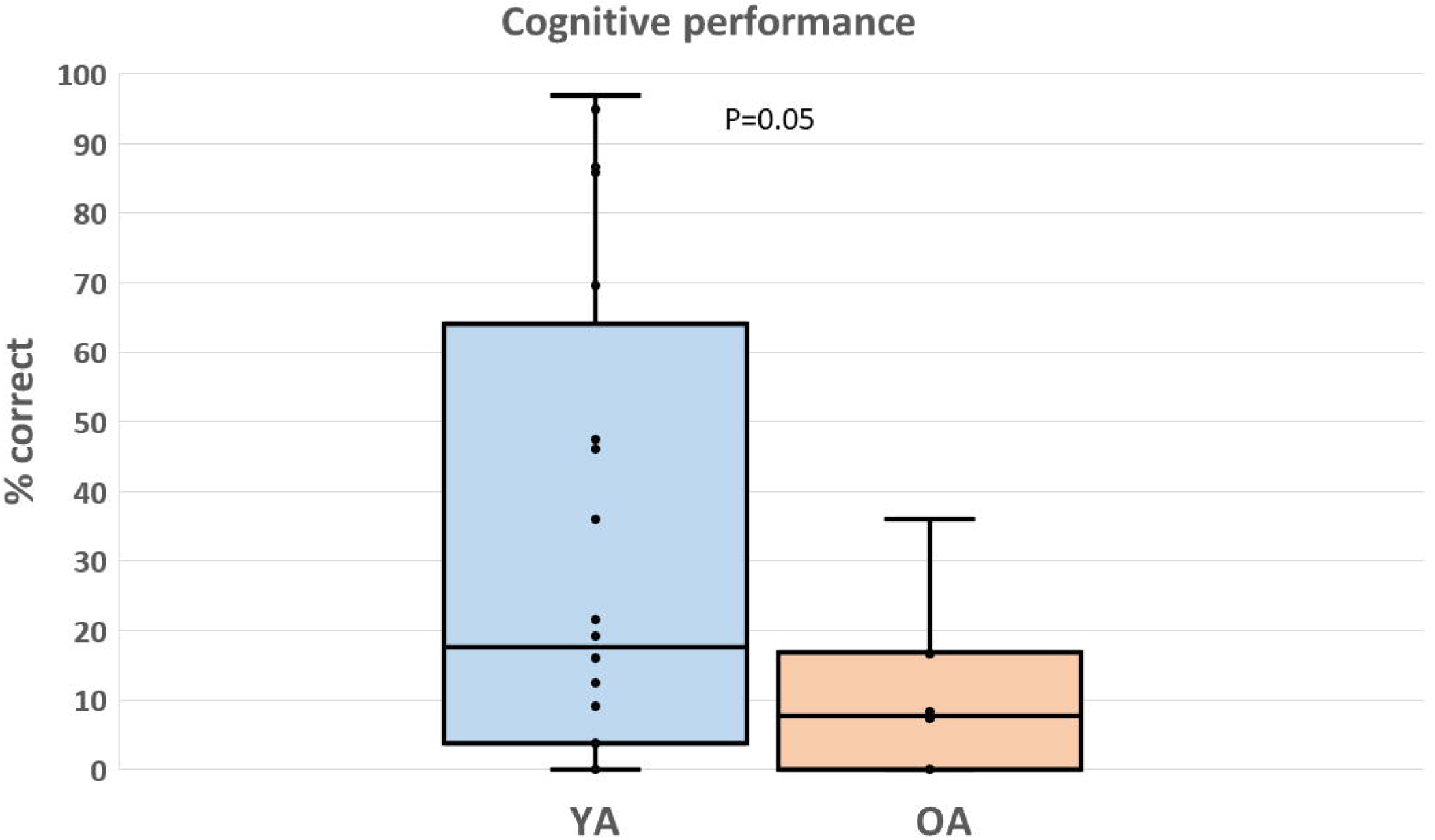
Percent of correct answers (Equation 2) in the concurrent cognitive dual task during perturbed-walking between young (n=19) and older adults (n=10) using the Mann-Whitney U-test. The median correct answers for young adults (19%) were significantly higher than the older adults (8%). Circles represent individual participants’ performance. Horizontal black lines represent group medians. Top and bottom of the boxplot represent the 1^st^ and 3^rd^ quartiles, respectively and whiskers represent the minimum and maximum values.

### Balance Recovery Performance

All participants successfully recovered from all perturbation trials, thus no falls occurred during the examinations. To achieve our aims, the data were split by perturbation type (i.e., the gait cycle phase during which the perturbation was implemented) and age groups– 108 perturbations in single-support (68 and 40 perturbations for YA and OA, respectively) and 152 perturbations in double-support (80 and 72 perturbations for YA and OA, respectively).

#### Effect of age on muscles activation pattern in response to perturbations during walking

In single support phase of gait cycle, YA showed a significant change in total spectral power for Left-VL muscle (adjusted p<0.001). OA showed a significant change in total spectral power for Left-TA muscle (adjusted p=0.011). Both groups demonstrated a significant change in total spectral power for Right-VL muscle (adjusted p≤0.011, Table 2).

**Table 2:**
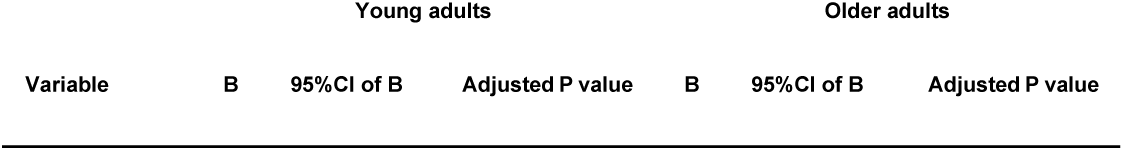
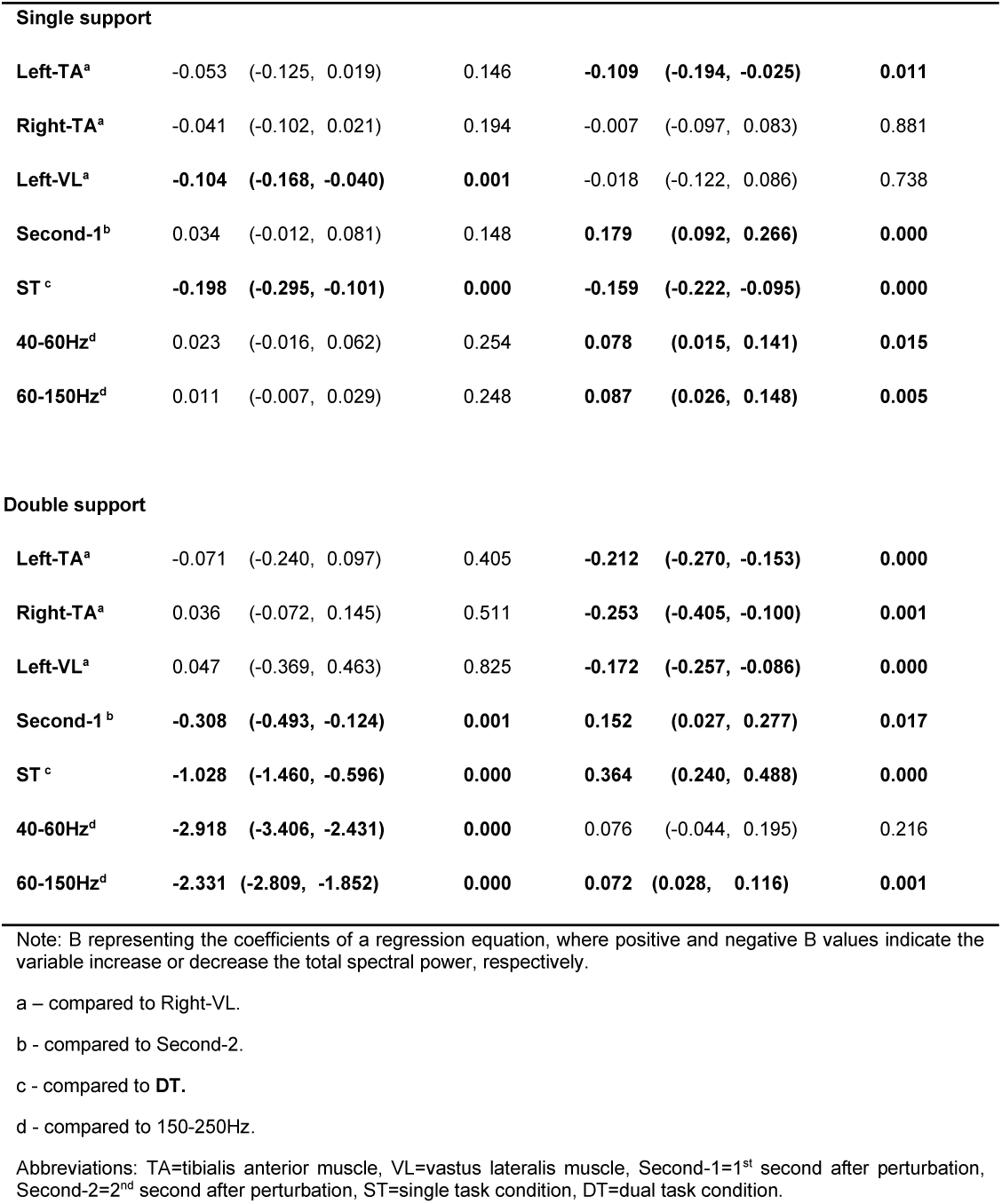
GEE models’ summary for muscle, time, cognitive load and frequency band effect on total spectral power in single-and double-support perturbations for young and older adults.

| Variable | Young adults |  |  | Older adults |  |  |
| --- | --- | --- | --- | --- | --- | --- |
|  | B | 95%CI of B | Adjusted P value | B | 95%CI of B | Adjusted P value |

| Single support |  |  |  |  |  |  |
| --- | --- | --- | --- | --- | --- | --- |
| Left-TA<sup>a</sup> | -0.053 | (-0.125, 0.019) | 0.146 | **-0.109** | **(-0.194, -0.025)** | **0.011** |
| Right-TA<sup>a</sup> | -0.041 | (-0.102, 0.021) | 0.194 | -0.007 | (-0.097, 0.083) | 0.881 |
| Left-VL<sup>a</sup> | **-0.104** | **(-0.168, -0.040)** | **0.001** | -0.018 | (-0.122, 0.086) | 0.738 |
| Second-1<sup>b</sup> | 0.034 | (-0.012, 0.081) | 0.148 | **0.179** | **(0.092, 0.266)** | **0.000** |
| ST<sup>c</sup> | **-0.198** | **(-0.295, -0.101)** | **0.000** | **-0.159** | **(-0.222, -0.095)** | **0.000** |
| 40-60Hz<sup>d</sup> | 0.023 | (-0.016, 0.062) | 0.254 | **0.078** | **(0.015, 0.141)** | **0.015** |
| 60-150Hz<sup>d</sup> | 0.011 | (-0.007, 0.029) | 0.248 | **0.087** | **(0.026, 0.148)** | **0.005** |
| Double support |  |  |  |  |  |  |
| Left-TA<sup>a</sup> | -0.071 | (-0.240, 0.097) | 0.405 | **-0.212** | **(-0.270, -0.153)** | **0.000** |
| Right-TA<sup>a</sup> | 0.036 | (-0.072, 0.145) | 0.511 | **-0.253** | **(-0.405, -0.100)** | **0.001** |
| Left-VL<sup>a</sup> | 0.047 | (-0.369, 0.463) | 0.825 | **-0.172** | **(-0.257, -0.086)** | **0.000** |
| Second-1<sup>b</sup> | **-0.308** | **(-0.493, -0.124)** | **0.001** | **0.152** | **(0.027, 0.277)** | **0.017** |
| ST<sup>c</sup> | **-1.028** | **(-1.460, -0.596)** | **0.000** | **0.364** | **(0.240, 0.488)** | **0.000** |
| 40-60Hz<sup>d</sup> | **-2.918** | **(-3.406, -2.431)** | **0.000** | 0.076 | (-0.044, 0.195) | 0.216 |
| 60-150Hz<sup>d</sup> | **-2.331** | **(-2.809, -1.852)** | **0.000** | **0.072** | **(0.028, 0.116)** | **0.001** |

Note: B representing the coefficients of a regression equation, where positive and negative B values indicate the variable increase or decrease the total spectral power, respectively.
a – compared to Right-VL.
b - compared to Second-2.
c - compared to DT.
d - compared to 150-250Hz.
Abbreviations: TA=tibialis anterior muscle, VL=vastus lateralis muscle, Second-1=1<sup>st</sup> second after perturbation, Second-2=2<sup>nd</sup> second after perturbation, ST=single task condition, DT=dual task condition.

In Double support phase of gait cycle, OA had significant association for lower limb muscles with the change in total spectral power, with the lowest change for Right-TA (adjusted p=0.001) and the highest for Right-VL (adjusted p≤0.001, Table 2) while YA did not show a muscle effect.

#### Age effect on muscle total spectral power in the first two seconds after perturbation onset

In single support phase of gait cycle, OA, showed a significant increase in EMG total spectral power in the 1^st^ Second compared to the 2^nd^ Second (adjusted p<0.001) while YA did not. In double support phase of gait cycle, OA had a greater increase in change in the 1^st^ Second compared to the 2^nd^ Second (adjusted p=0.017). However, YA showed greater increase for the 2^nd^ Second compared to the 1^st^ second (adjusted p=0.001, Table 2).

#### Age-related effects on muscle fiber type recruitment in response to unexpected perturbations during walking

In single support phase of gait cycle, OA had the greatest change in total spectral power for 60-150 Hz, type IIa related frequencies (adjusted p=0.015) and the lowest for 150-250Hz, Type IIb related frequencies (adjusted p=0.005) while YA did not show an effect of frequency band. In double support phase of gait cycle, OA had significantly greater change in 60-150Hz, type IIa related frequencies, compared to 150-250Hz, type IIb related frequencies (adjusted p=0.001). YAs had the greatest change for 150-250Hz, type IIb related frequencies (adjusted p<0.001) and the lowest for 40-60Hz, Type I related frequencies (adjusted p<0.001, Table 2).

#### The effect of concurrent cognitive task on the total EMG spectral power in response to perturbation during walking

YA and OA show a greater change in the total spectral power in response to perturbed DT condition compared to perturbed ST condition in single support phase (adjusted p<0.001). In double support phase perturbations, YA presented a similar pattern to the single support phase perturbations where perturbed DT condition had greater increase in total spectral power compared to perturbed ST condition (adjusted p<0.001). OA however, showed greater increased total spectral power in perturbed ST condition compared with perturbed DT condition in double support phase (adjusted p<0.001, Table 2).

#### Muscles activation and the association with DTC

For YA, we found that during perturbed DT and ST conditions the greatest increase in DTC was associated with Right-TA (adjusted p=0.012, adjusted p=0.028, respectively) followed by Left-TA (adjusted p=0.018, adjusted p=0.054, respectively). OA had no significant muscle effect (adjusted p≥0.110) in single support phase of gait cycle and lowest increase of DTC association with Right-TA (adjusted p=0.002), followed by Left-VL (adjusted p=0.012) and Left-TA (adjusted p=0.066), in perturbations during the double support phase of gait cycle (Table 3).

**Table 3:**
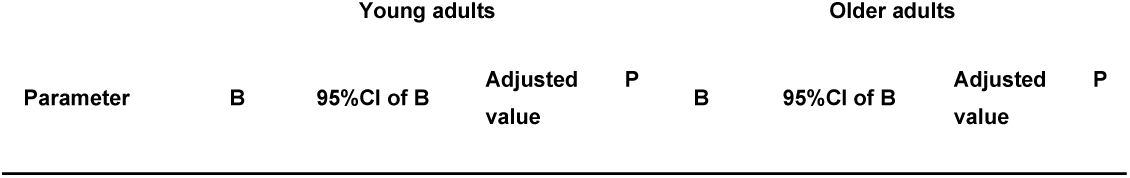
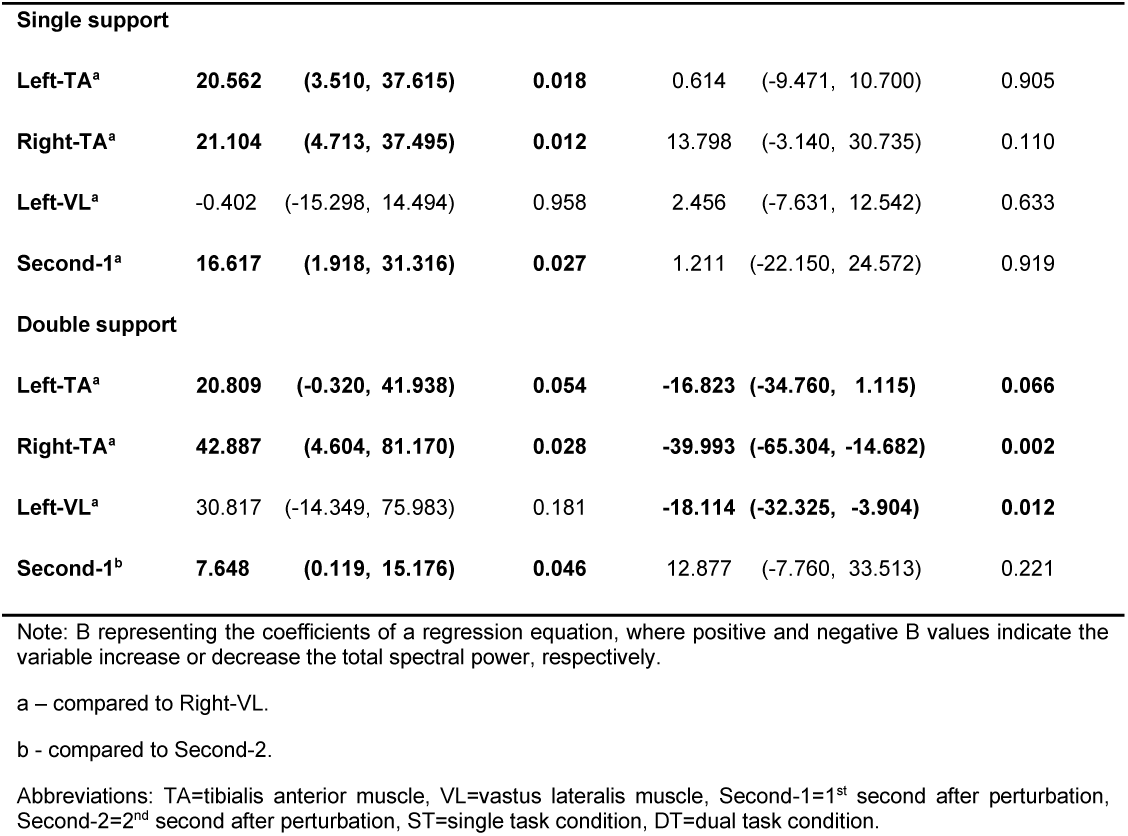
GEE models’ summary for muscle and time effects on dual task cost in single-and double-support perturbations for young and older adults.

#### Association between the time window and DTC

For YA, in perturbed DT and ST conditions, DTC increase had greater association with 1^st^ Second compared to 2^nd^ Second (adjusted p=0.027, and adjusted p=0.046, respectively). For OAs, no significant effect of time was found on DTC after perturbations in single or double support phases of gait (Table 3).

Figure 1S, in *supplementary materials*, show graphical representation of DTCs of the different muscles in 1^st^ Second and 2^nd^ Second and Figure 2S, in *supplementary materials*, for DTCs correlation analysis for exploring age effect on the associations between muscles’ recruitment patterns in response to unexpected perturbations during walking.

## Discussion

Previous works have studied the differences in muscle activation magnitude between young and older adults in response to unexpected perturbations during standing (e.g., [47]) and walking (e.g., [48]). To the best of our knowledge, the effects of age and a concurrent cognitive task on muscle fiber recruitment patterns in response to unannounced perturbations during walking was not explored. We used wavelet analysis to study muscle fiber related frequencies in the sEMG content of YA and OA. We found a dominant increase in type IIa muscle fiber type related frequencies (60-150Hz) for OA, in single-and double-support phases of gait perturbations compared to the greatest increase in type IIb muscle fiber type related frequencies (150-250Hz) for YA in double-support phase of gait, supporting our first hypothesis that age will affect muscle type recruitment. In addition, we found that YA show a greater total spectral power increase in response to perturbations during DT condition compared to ST condition, regardless of the perturbation type. OA, however, showed different patterns across perturbation types, suggesting a difference in motor task demand. These partially support our second hypothesis. Also, dividing the data into single-and double-support phases of gait perturbations provided more information than averaging all perturbation conditions.

### Cognitive performance

#### Age effect on concurrent cognitive task performance in perturbed gait

Our results agree with previous investigations, showing deterioration in concurrent cognitive task performance with age[49]. Rapp et al. [49] combined a working memory task with a postural control task. We show this holds for combined working memory (i.e., paced auditory serial addition task) with perturbations during walking.

### Balance reactive performance

#### Muscles engaged in the response to unexpected perturbations differ with age and show different effect on DTC

As a result of an unannounced surface translation perturbation while walking, there is a change in the relationship between the CoM and the base of support (i.e., center of pressure (CoP); [50]. It is well-established that to control these variables, a person utilizes the ankle strategy (i.e., activation of ankle muscles in response to balance perturbations) to shift the CoP in relationship to the moving CoM. When this strategy is no longer effective, a step in the direction of the CoM displacement is executed[51]. This postural balance strategy is mediated by the ankle muscles (e.g., TA), effectively leading to better control of CoP displacement in YA, than OA. Similar results were found in other studies (e.g., [52]).

Results of our GEE models show that: (1) both YA and OA increase activation of the thigh muscle of the reacting limb (i.e., Right-VL) the most in response to perturbations during walking. Our results are in agreement with those of Tang and Woollacott [48] who showed greater burst magnitude for the anterior thigh muscle (rectus femoris muscle) than the anterior leg muscle (tibialis anterior) of the reacting limb in response to slip perturbations during walking, both for YA and OA. Our results suggest that the anterior thigh muscles play a crucial role in balance control during perturbed walking, regardless of perturbation direction, probably to control the center of mass (CoM) vertical movement, i.e., working against gravity. (2) Regarding the supporting limb (i.e., left lower limb) YA relies less on VL while OA relies less on TA. Tang and Woollacott [48] did not find significant differences between the muscles, possibly due to the difference in perturbation direction implemented in their study. Our results suggest that muscle activations of the supporting limb are more affected by perturbation direction or mechanism compared to the reacting limb. It also suggests that YA try to implement the ankle strategy (i.e., activation of leg muscles to control the ankle CoM displacement in response to perturbations) while OA, are less likely to apply this strategy. Mackey and Robinovitch [47] found that in response to tether release perturbations during standing (i.e., leaning forward supported and losing the support unexpectedly), OA women exhibit small ankle muscles peak torque, slower reaction times and slower ankle torque generation, explaining differences in balance recovery performance with the ankle strategy between YA and OA women (male were not tested in that study). Furthermore, [53] found similar results for walking, where hip and knee concentric powers during the stance phase of walking were higher in OA than in YA despite their decreased force generation of the ankle muscles. [54] found that, across locomotion modes of walking, running, and sprinting, older men reduced the ankle extensor force generation. Finally, our second set of GEE models supports these findings by showing that the increased DTC has the strongest association with Right-TA for YA and Right-VL for OA. This implies that YA and OA employ different strategies to keep their balance while responding to walking perturbations during DT condition.

#### YA and OA muscle activations, concurrent cognitive task and DTC have different associations with time after perturbation onset

To the best of our knowledge, we are the first to show the dynamics of muscle activation pattern differences between YA and OA over time after unexpected perturbations. We found that OA consistently show higher increase in Second-1 compared to Second-2.

Furthermore we observed age differences in the increase in DTC. YA show consistently greater increase in Second-1 compared to Second-2 (i.e., in DTC), while OA show no significant effect of time. We therefore propose that OA are more affected by the perturbed DT condition as compared to YA i.e., the effect lasts longer than in YA, suggesting the OA ‘stay alert’ for longer, in the ‘recovery process’ from a perturbation when exposed to secondary cognitive task.

This also suggests that the recovery process from perturbations requires cognitive resources. A DT paradigm involves concurrently performing both attention-demanding and balance tasks.

While the literature has not addressed the dynamics of DTC after perturbations, findings from studies using a DT paradigm showed a decrease in performance of postural tasks, or cognitive tasks, or both among young adults, as well as older adults [55–59]. In some cases, DT improved postural stability[60,61]. The inconsistencies in the results depend on the postural and cognitive challenges[62]. Thus, the extent of the interference seems to depend on motor demands, central processing abilities, and attentional costs[63,64].

The assumption is that resource competition occur since the available central processing resources are assumed to be limited[64]. This assumption is supported by the positive DTC we found. In a systematic review, Wittenberg et al.[16] provide evidence to support an increase in brain activation in balance control tasks, regardless of mechanical, cognitive, or sensory challenges. Thus, DT performance may indicate the involvement of high cortical brain structures, such as the supplementary motor area, in these tasks.

#### Concurrent cognitive task affects total spectral power in response to physical perturbations during walking

We found a significant increase in total spectral power in response to perturbations with a concurrent cognitive task in both age groups. This result agrees with [65], which found elevated muscle activation of bilateral leg and thigh muscles in response to perturbations during walking. It also corresponds with literature providing evidence of DT interference in postural performance amongst YA[66]. Little and Woollacott[66] showed competition for central processing attentional resources between the postural control and concurrent cognitive DT, through the reduction in amplitude of event-related potentials over motor and sensory cortical areas amongst YA. They found no DTCs on EMG activity. The disagreement in the findings of EMG DTC may arise from the implementation of different perturbation paradigms. While Little and Woollacott[66] perturbed their participants in standing, we perturbed our participants in walking, which may be more demanding and ecologically valid. Also, we examined the spectral power content while Little and Woollacott examine the EMG magnitude and latency. Lastly, a systematic review by Boisgontier et al.,[67] showed that when the complexity of a postural task is increased by dynamic conditions (e.g., support surface perturbations), performance in postural, cognitive, or both tasks is more affected in OA relative to YA.

#### Age affects muscle fiber type recruitment in response to unexpected perturbations during walking

The results of the GGE models show greatest association of type IIa associated frequencies (60-150Hz) with change in total spectral power for OA in both single-and double-support perturbations. YA show greater association of type IIb associated frequencies (150-250Hz) with change in total spectral power in double-support perturbations only. These results suggest that the ability to recruit fast-twitch type IIb muscle fibers is deteriorated with age. Supporting results were introduced in the sports and exercise literature by [68]. They found that sprinters with a higher percentage of thigh and calf circumference and skeletal muscle mass, have a better capability to recruit fast twitch muscle fibers. Although we did not measure thigh and calf circumference and skeletal muscle mass, loss of muscle mass with aging, is associated with reduced type IIb muscle fiber number and size, and very common with aging [69].

## Limitations

Findings described herein should be considered in the context of several limitations. First, the DT implemented was an arithmetic paced serial addition task, where the participants had to add 15 numbers that were read to them at a pace of one number per two seconds and provide the sum at the end of the reading. As a result, only accuracy rate could be inferred. Some of the participants in our age groups had 0% correct answers (see Figure 2). This raises a concern for their engagement or understanding of the DT condition. We found a positive DTC in the physiological data of all participants (see Figure 1S in the *Supplementary materials*) suggesting engagement with the task even when they failed to perform the DT correctly.

Another limitation is the relatively weak (in terms of magnitude of acceleration) set of perturbations employed in our protocol compared, for example, to [70,71]. It is therefore possible that our results provide a partial picture regarding potential muscle activation pattern in response to perturbations in a larger magnetite spectrum. Also, it should be considered that this study examined anterior lower limb muscles. Future studies including additional muscles (e.g., Glutei, Hamstrings, Lower-back and Upper-limbs muscles) and higher perturbation magnitudes would provide a more complete picture of muscle activation patterns in response to unexpected perturbations during walking. Lastly, although our methodology is consistent with the extensive existing literature implementing sEMG (see Methods), signal reflecting ‘cross-talks’ cannot be ruled out. In this context, VL is a deeper muscle relative to TA, making it more susceptible to cross-talks.

## Acknowledgments

This study was supported in part by funding from the Israeli Ministry of Science and Technology, grant #3-12072, and from the Israel Science Fund, grant #3-14527. The research is part of one contributor’s (UR) work towards a doctoral degree from Ben-Gurion University of the Negev and was partially supported by a stipend.

## Funding and/or Conflicts of interests/Competing interests

The authors declare no conflict of interests.

## Notes

### Competing Interest Statement

The authors have declared no competing interest.

