## Supplementary Materials for "Age and cognitive load affect muscle activation profiles in response to physical perturbations while walking"

### Methods

To test the relationship between the DTCs of muscles' activation, for the different muscles, for both YA and OA we used the median DTC value for every participant in a spearman correlation, since the medians did not distribute normally.

### Results

#### *Age effect on the associations between muscles' recruitment patterns in response to unexpected perturbations during walking*

To investigate the age effect on muscles' recruitment patterns in response to lateral perturbations during walking we calculated the correlation between the DTCs for the different muscles in Second-1 and Second-2 for the different age groups. Data are presented in Figure 1S and the correlation results are summarized in Figure 2S.

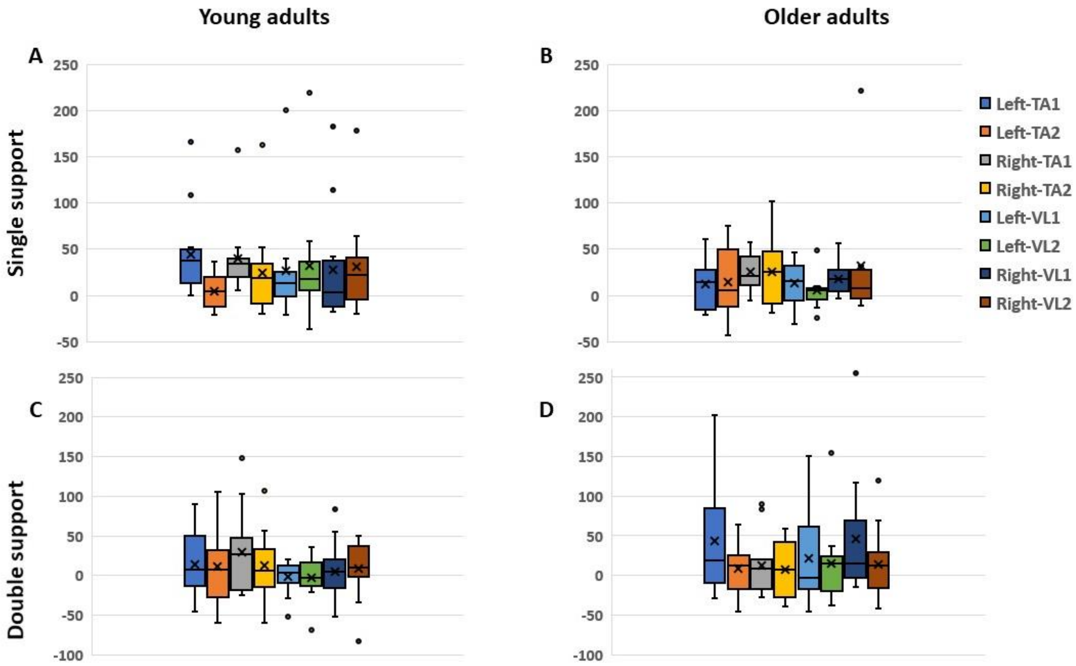


**Figure 1S: Boxplots of dual task cost (DTC).** Depicting DTC for walking with perturbations while performing a concurrent cognitive task in single support for young (A) and older adults (B), and in double support (C & D, respectively). Boxplots borders represent the 1st and 3rd quartiles. Horizontal line represents the median, while X represents the mean. Whiskers represent minimum and maximum values.

#### *Single support*

*Young adults*

DTC for Left-TA in Second-1 is in positive high correlation with DTC for Right-VL activity in Second-2 ($r=0.72$, $p<0.05$). Left- and Right-VL DTCs in Second-1 are in moderate and high positive correlation with DTC for Left-TA in Second-2 ($r=0.66$ and 0.76, $p<0.05$, respectively). DTC for Left-TA in Second-2 is in positive high correlation with DTC for Right-TA and Right-VL in Second-2 ($r=0.85$ and $r=0.74$, $p<0.01$, respectively). DTC for Right-TA in Second-1 is in positive moderate correlation with DTC for Right-VL in Second-2 ($r=0.63$, $p<0.05$). DTC for Right-TA in Second-2 is in positive high correlation with DTC for Right-VL in Second-2 ($r=0.81$, $p<0.01$). DTC for Left-VL in Second-1 is in positive high correlation with DTC for Right-VL in Second-1 ($r=0.71$, $p<0.05$. Finally, DTC for Right-VL in Second-1 is in positive moderate correlation with DTC for Right-VL in Second-2 ($r=0.63$, $p<0.05$). See Figure 2S, panel A.

*Older adults*

DTC for Left-TA in Second-1 is in negative high correlation with Right-TA in Second-2 ($r=-0.79$, $p<0.05$). DTC for Left-TA in Second-2 is in high positive correlation with Left-VL in Second-2 ($r=0.86$, $p<0.01$). Finally, DTC for Right-TA in Second-1 is in high positive correlation with Right-VL in Second-1 ($r=0.76$, $p<0.05$). See Figure 2S, panel B.

***Double support***

*Young adults*

DTC for Left-TA in Second-1 is in moderate to high correlation with Left-TA, Right-TA, Left-VL, and Right-VL in Second-2 ($r=0.81$, $r=0.57$, $r=0.60$, $r=0.67$, $p<0.05$, respectively). DTC for Left- and Right-TA in Second-2 is moderately to highly correlated with all muscles in Second-1 and Second-2 ($0.90\geq r\geq 0.53$, $p<0.05$). DTC for Right-TA in Second-1 is in positive moderate to high correlation with Left-VL, and Right-VL in Second-1 ($r=0.77$ and $r=0.64$, $p<0.05$, respectively) and highly correlated with Left- and Right-TA and Left-VL in Second-2 ($r=0.80$, $r=0.88$, and $r=0.71$, $p<0.01$). DTC for Left-VL in Second-1 is in moderate correlation with Left-VL in Second-2 ($r=0.61$, $p<0.05$). Finally, DTC for Left-VL in Second-2 is in positive moderate correlation with Right-VL in Second-2 ($r=0.62$, $p<0.62$). See Figure 2S, panel C.

*Older adults*

DTC for Left-TA in Second-1 is in positive high correlation with Right-TA and Left-VL in Second-1 (r=0.72 and r=0.76, p<0.01, respectively). DTC for Left-TA in Second-2 is in positive high correlation with Left- and Right-VL in Second-2 (r=0.74 and r=0.89, p<0.01). DTC for Right-TA in Second-1 is in positive moderate correlation with Right-TA in Second-2 (r=0.59, p<0.05). DTC for Left-VL in Second-1 is in positive moderate correlation with Right-VL in Second-1 (r=0.69, p<0.05). Finally, DTC for Left-VL in Second-2 is in positive high correlation with Right-VL in Second-2 (r=0.76, p<0.01). See Figure 2S, panel D.

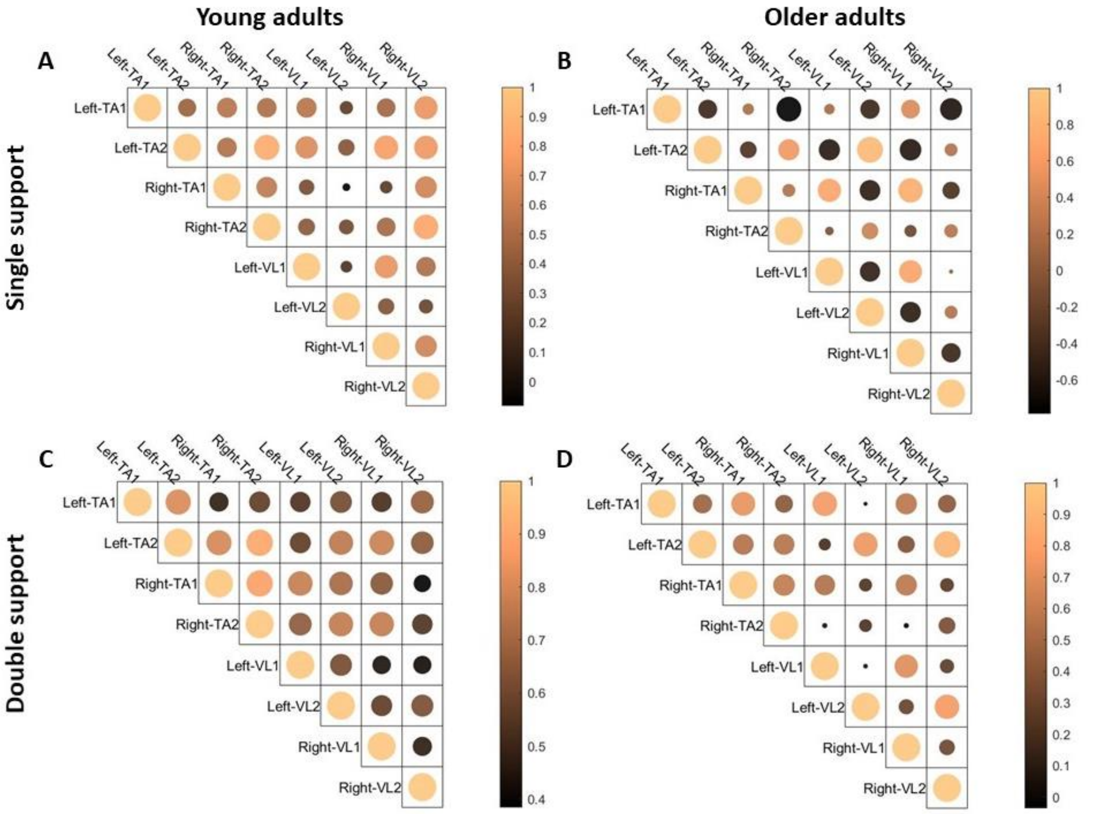


**Figure 2S: Correlation matrices for tibialis anterior (TA) and vastus lateral (VL) muscles' dual task costs in Second-1 and Second-2.** Correlations in response to perturbations in single support condition for young and older adults are presented in panels (A) and (B), respectively. Double support condition correlations are depicted in panels (C), young adults, and (D), older adults. Colors and circles' size represent correlation strength, with larger circles representing higher, negative or positive, correlations. Notice the different scales and color codes for the different panels. Values of r > 0.6 were found significant, see main text for specific p values.

### Discussion

#### *Age effect on the associations between muscles' DTC in response to unexpected perturbations during walking*

The results of the correlation analysis for single and double support perturbations YA express more significant correlations in DTC across and within muscles compared to OA. This shows a more synchronized response between the leg and the thigh muscles in YA compared to OA. The higher synchrony persists from Second-1 to Second-2. Our results support previous study by Tang et al.[1] who found synchronized activity from bilateral leg and thigh muscles and the coordination between the two lower limbs were the essential component to reactive balance control in YA. It appears that OA lack some of the synchronous muscle activation which might be affecting their reactive balance control.

Our correlation analysis revealed more synchronized muscle response for YA compared to OA.
